# Checkpoint-independent association of Mad3^BUBR1^ with Stu1^CLASP^ directs homolog alignment in meiosis I

**DOI:** 10.1101/2024.01.04.574163

**Authors:** Anuradha Mukherjee, Christos Spanos, Adele L. Marston

## Abstract

Gametes are produced via meiosis, a specialized cell division associated with frequent errors which cause birth defects and infertility. Uniquely in meiosis I, homologous chromosomes segregate to opposite poles, usually requiring their linkage by chiasmata, the products of crossover recombination^1^. The spindle checkpoint delays cell cycle progression until all chromosomes are properly attached to microtubules^2^ but the steps leading to the capture and alignment of chromosomes on the meiosis I spindle remain poorly understood. In budding yeast meiosis I, Mad2 and Mad3^BUBR1^ are equally important for spindle checkpoint delay, but biorientation of homologs on the meiosis I spindle requires Mad2, but not Mad3^BUBR1^ ^3,4^. Here we show that Mad3^BUBR1^ promotes accurate meiosis I homolog segregation outside its canonical checkpoint role, independently of Mad2. We find that Mad3^BUBR1^ associates with the TOGL1 domain of Stu1^CLASP^, a conserved plus-end microtubule protein which is important for chromosome capture onto the spindle. Homologous chromosome pairs that are proficient in crossover formation, but which fail to biorient, rely on Mad3^BUBR1^-Stu1^CLASP^ to ensure their efficient attachment to microtubules and segregation during meiosis I. Furthermore, we show that Mad3^BUBR1-^Stu1^CLASP^ are essential to rescue the segregation of mini-chromosomes lacking crossovers. Our findings define a new pathway ensuring microtubule-dependent chromosome capture and demonstrate that spindle checkpoint proteins safeguard the fidelity of chromosome segregation both by actively promoting chromosome alignment and delaying cell cycle progression until this has occurred.

## Results and discussion

### Distinct, checkpoint-independent, functions for Mad2 and Mad3^BUBR1^ in homolog segregation during meiosis I

Meiosis I cell cycle delay in response to either unattached kinetochores or a lack of inter-homolog tension requires Mad2 and Mad3^BUBR1^, indicating that both proteins are essential for a functional spindle checkpoint^3,4^. In contrast, in an unperturbed meiosis, Mad2 is more important than Mad3^BUBR1^ for chromosome segregation, since Mad2, but not Mad3^BUBR1^, is required for homolog biorientation^4^ (Figure S1A and B). Together, these findings indicate that Mad2 promotes homolog biorientation and chromosome segregation independently of the spindle checkpoint-mediated delay. Furthermore, we observed additive effects of *mad2Δ* and *mad3Δ* on the non-disjunction of homologs during meiosis I. Cells with both homologs of chromosome V labelled close to the centromere (*CEN5-tdTomato*) and carrying a spindle marker (*GFP-TUB1*) were induced to enter meiosis and followed by live cell imaging (Figure 1A-C). As described previously^4^, homolog non-disjunction in meiosis I was only mildly elevated in *mad3Δ* (<5%) compared to wild type cells, while ~10-15% of *mad2Δ* cells failed to segregate homologs to opposite poles in meiosis I (Figure 1C). However, deletion of *MAD3* in the *mad2Δ* background increased meiosis I non-disjunction to ~20% (Figure 1C). Given that both *mad2Δ* and *mad3Δ* lack a functional spindle checkpoint^3^, their additive effects reveal distinct spindle checkpoint-independent functions of Mad2 and Mad3^BUBR1^ in meiosis I chromosome segregation (Figure 1D). Importantly, Mad3^BUBR1^ plays only a minor role on homolog segregation in otherwise wild type cells but becomes critical in the absence of Mad2, suggesting that Mad3^BUBR1^ provides a backup mechanism when biorientation is compromised.

**Figure 1.**
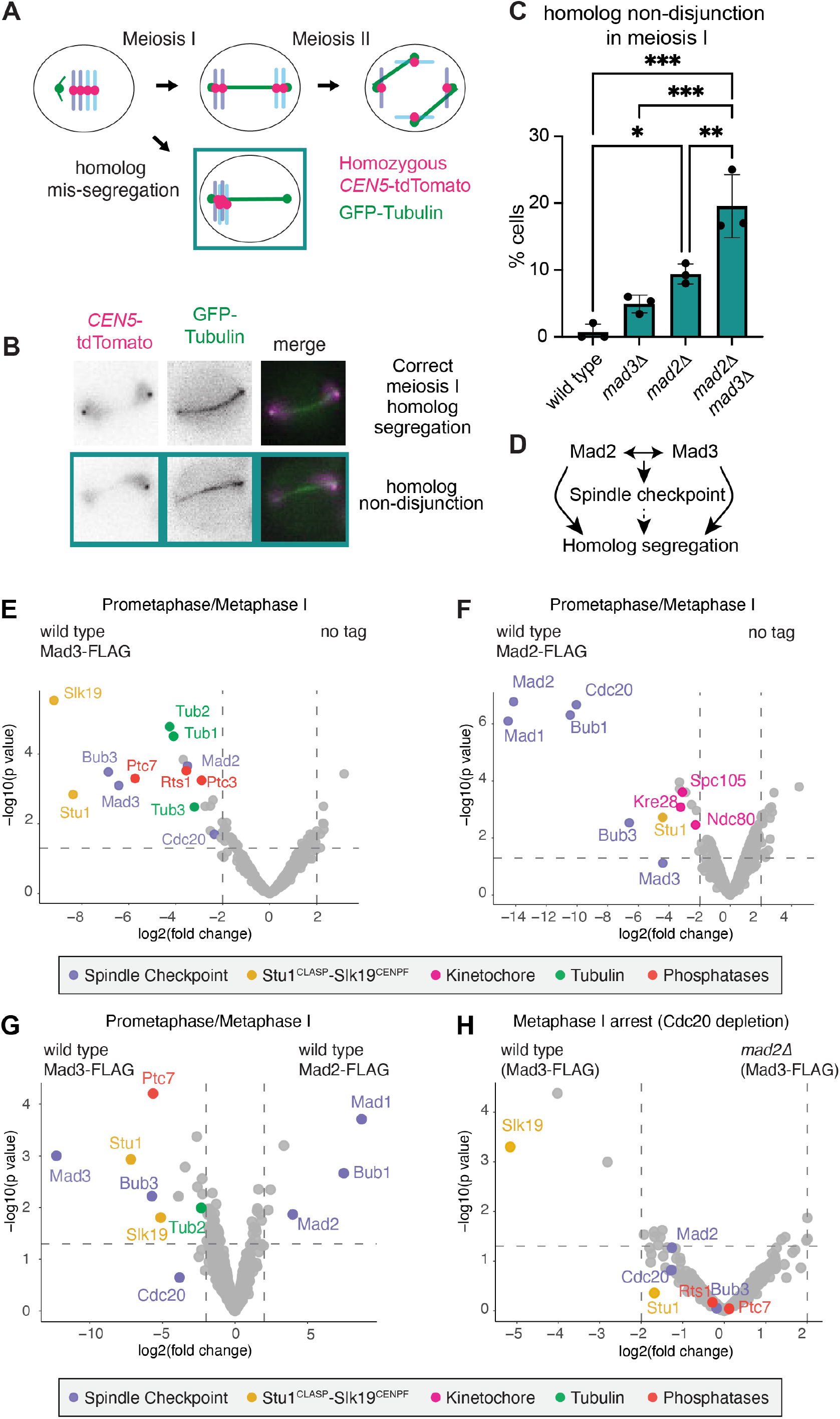
Mad2 and Mad3^BUBR1^ have distinct, spindle-checkpoint independent functions in meiosis I. (A-C) Mad2 and Mad3 show additive effects on homolog segregation in meiosis I. Cells carrying *CEN5*-GFP and *GFP-TUB1* were induced to sporulate and live imaged. (A) Schematic of the experiment. (B) Representative images of correct meiosis I segregation and non-disjunction. (C) Percentage of cells of the indicated genotypes showing meiosis I non-disjunction. Mean of three biological repeats (*n*=21-60, wild type; *n*=56-68 *mad31*; *n*=53-65 *mad21*; *n*=57-60 *mad21 mad31*). Error bars represent standard deviation, **** p<0.0001, ***p≤0.001, **p≤0.01, *p≤0.05, One way ANNOVA (Tukey’s multiple comparisons test). (D) Summary depicting the shared roles of Mad2 and Mad3 in the spindle checkpoint and separate roles in directing homolog segregation in meiosis I. (E-G) Immunoprecipitation and mass spectrometry of Mad2-FLAG and Mad3-FLAG during prometaphase/metaphase I. Volcano plots showing the relative enrichment of proteins immunoprecipitated with (E) Mad3-FLAG compared to no tag, (F) Mad2-FLAG compared to no tag conditions and (G) Mad3-FLAG vs. Mad2-FLAG. Cells were harvested 75 min after release from a prophase I arrest (corresponding to prometaphase/metaphase I). (H) Mad3-FLAG interacts with Stu1 independently of the spindle checkpoint. Volcano plot showing the comparative enrichment of proteins identified by mass spectrometry in Mad3-FLAG immunoprecipitates from wild type and *mad21* cells harvested 6h after inducing sporulation where progression beyond metaphase I was prevented by depletion of Cdc20 (*pCLB2-CDC20*). Results in E-G include data from three biological replicates, and H includes data from two biological repeats for each condition. Log_2_(Fold Change) between conditions is shown with corresponding p values. Dashed line indicates Log_2_(Fold Change) = |2|.

### Mad3^BUBR1^ specifically interacts with Stu1^CLASP^, independently of the spindle checkpoint

To uncover the role of Mad3^BubR1^ in homolog segregation we sought to identify interacting proteins in meiosis I. Wild-type cells carrying FLAG-tagged Mad3^BUBR1^ or Mad2 and a no tag control were harvested 75 min after release from a prophase block (corresponding to prometaphase/metaphase I (Figure S2A)) and anti-FLAG immunoprecipitates were analysed by mass spectrometry. This revealed interacting proteins common to both Mad2 and Mad3^BUBR1^ but also some proteins specific to each (Figure 1E-G). Spindle checkpoint proteins were found in both cases, with Mad2 predominantly binding Mad1, Bub1 and Cdc20, while Bub3 is the major Mad3^BUBR1^ interactor, consistent with known direct binding events^5–8^. However, proteins of the outer kinetochore, including Ndc80 and Spc105^KNL1^-Kre28, to which Bub1 binds directly as part of its role in the spindle checkpoint^9^, were found only in Mad2 immunoprecipitates (Figure 1F and G). In contrast, several distinct proteins, including tubulin subunits (Tub1, Tub2 and Tub3), three distinct phosphatases (Ptc3, Ptc7 and Rts1, a subunit of PP2A-B56) and the microtubule-regulator Stu1^CLASP^, along with its binding partner Slk19^CENPF^ were significantly enriched in the Mad3^BUBR1^ purification (Figure 1E and G). Direct comparison confirmed that Stu1^CLASP^, Slk19^CENPF^ and Ptc7, in addition to Bub3, were most significantly enriched in Mad3^BUBR1^ over Mad2 purifications (Figure 1G). Since Mad3^BUBR1^ promotes homolog segregation independently of the spindle checkpoint or Mad2 (Figure 1A-C), we reasoned that relevant interactions in this process should persist in cells where the spindle checkpoint has been abrogated. To test this idea, we used mass spectrometry to compare Mad3-FLAG immunoprecipitates with or without Mad2 (to abrogate the checkpoint) in Cdc20-depleted cells (metaphase I arrest). Interestingly, while the association of Slk19^CENPF^ with Mad3^BUBR1^ was greatly diminished in *mad2Δ* cells, the interactions with Stu1^CLASP^ and Ptc7 were maintained (Figure 1H). Stu1^CLASP^, but not Slk19^CENPF^, was also enriched to some extent in Mad3 ^BUBR1^ immunoprecipitates from meiotic prophase I-arrested cells (Figure S2B). We conclude that Mad3^BubR1^ associates with Stu1^CLASP^ independently of the spindle checkpoint and Mad2.

### Mad3^BUBR1^ interacts with Stu1^CLASP^ through its TOGL1 domain

Stu1^CLASP^ is a member of the conserved CLASP family of microtubule regulators which suppress catastrophes and promote rescue of microtubule plus ends^10^. CLASPs are critical in cell migration, neuronal development and during mitosis, where they are essential for the capture and alignment of chromosomes on the mitotic spindle^10^. This makes Stu1^CLASP^ an attractive candidate for mediating the non-checkpoint functions of Mad3^BubR1^. Like all CLASP proteins, Stu1^CLASP^ is a modular protein with distinct functional domains (Figure 2A)^11^. We reasoned that the N-terminal TOGL1 domain, which is reported to be required for Stu1^CLASP^ localization at kinetochores but not its binding to microtubules or viability^11^, might be relevant for the Mad3^BUBR1^ interaction. To test this idea, we generated strains producing Stu1^CLASP^ lacking the TOGL1 domain by deletion of amino acids 17-260, hereafter called Stu1ΔTOGL1. To circumvent detrimental effects of expressing only Stu1ΔTOGL1 in mitosis, these cells also carried *STU1* under control of the mitosis-specific *CLB2* promoter (*pCLB2-STU1*), which is shut off upon meiotic entry resulting in production of Stu1ΔTOGL1 but not Stu1. Analysis of Mad3-FLAG immunoprecipitates in wild type and *stu1ΔTOGL1* cells by mass spectrometry indicated that Stu1^CLASP^, but not Stu1ΔTOGL1 co-purified with Mad3^BUBR1^ in both prometaphase I and prophase I cells (Figure 2B-D). Therefore, Mad3^BUBR1^ association with Stu1^CLASP^ requires its N-terminal TOGL1 domain.

**Figure 2.**
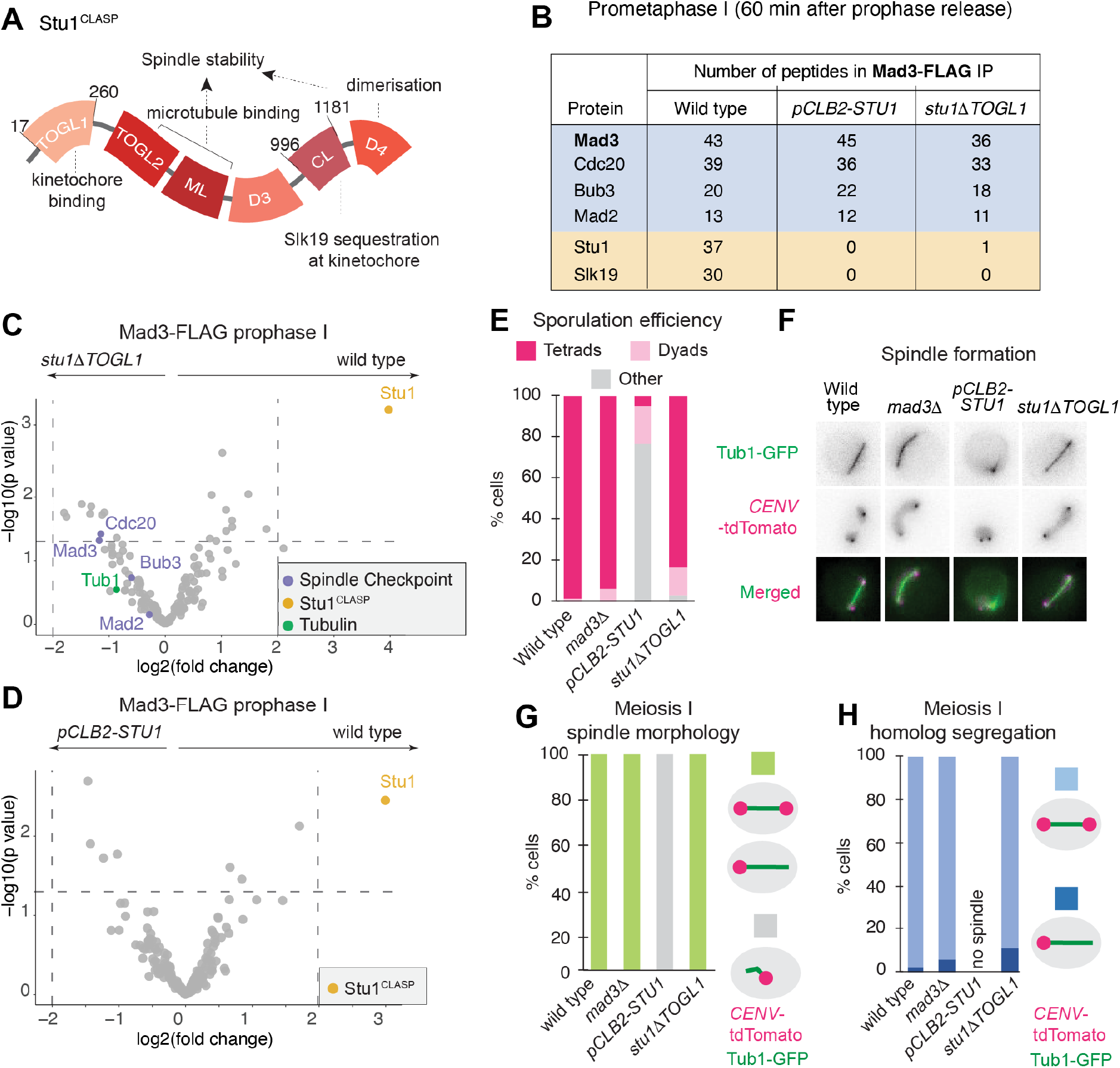
Mad3^BUBR1^ interacts with the TOGL1 domain of Stu1^CLASP^. (A) Schematic of Stu1 protein with domains shown as identified by^11^. (B) List of proteins and their unique peptide counts as identified by one repeat of mass spectrometry after immunoprecipitation of Mad3-FLAG in the indicated strains. Note that all three strains have heterozygous *pCLB2-STU1*, with the other allele as indicated. Cells were harvested 60 min after release from prophase I. (C and D) Mad3 interaction with Stu1 is lost in *stu11TOGL1* cells. Volcano plots after mass spectrometry showing the relative enrichment of proteins immunoprecipitated with Mad3-FLAG in (C) wild-type vs *stu11TOGL1* and (D) wild-type vs *pCLB2-STU1* prophase I-arrested cells. (E) Stu1, but not its TOGL1 domain, is required for sporulation. The percentages of tetrads and dyads produced 72 h after inducing sporulation was scored for 200 cells of the indicated genotypes. (F-H) Stu1, but not its TOGL1 domain, is required for spindle formation and chromosome segregation during meiosis I. Strains of the indicated genotypes and carrying *CEN5-tdTomato* and *GFP-TUB1* were induced to sporulate and live imaged. Representative images (F), together with the scoring of spindle morphology (G) and *CEN5-tdTomato* segregation (H) scored in anaphase I cells (*n*=54 wild type; *n*=50 *mad31*; *n*=50 *pCLB2-STU1*; *n*=56 *stu11togl1)*. Note that anaphase I spindles were not observed in *pCLB2-STU1* cells.

Stu1^CLASP^ plays a critical role in microtubule organisation and, accordingly, *STU1* is essential for the viability of mitotically cycling cells^12^. To understand whether Stu1^CLASP^ is similarly important in meiosis we analysed cells where both copies of *STU1* are under control of the *CLB2* promoter to prevent its expression in meiosis. Upon induction of sporulation, virtually all wild-type and *mad3Δ* cells underwent meiosis to produce four gametes, called a tetrad (Figure 2E). In contrast, over 75% of *pCLB2-STU1* cells failed to produce spores, with the remaining cells predominantly producing dyads (two spores), rather than tetrads. However, sporulation was largely unperturbed in *stu1ΔTOGL1* cells, with over 80% of cells forming tetrads (Figure 2E). Therefore, although Stu1 is critical for sporulation, its N-terminal TOGL1 domain, which is required for Mad3^BUBR1^ association, is not. Next, we used live-cell imaging to determine the requirement for Stu1^CLASP^, and its TOGL1 domain in spindle formation and chromosome segregation during meiosis. Cells carrying *GFP-TUB1* (to label microtubules) and homozygous *CEN5-tdTomato* to label both homologs of chromosome V were induced to sporulate and live imaged. Anaphase I spindles were observed in wild-type, *mad3Δ* and *stu1ΔTOGL1* cells but never in *pCLB2-STU1* cells, indicating that similar to mitosis^12^, Stu1 is required for bipolar spindle formation in meiosis I (Figure 2F and G). As a further confirmation of the specificity of the *stu1ΔTOGL1* allele we purified Stu1-GFP or Stu1ΔTOGL1-GFP and identified interacting proteins by mass spectrometry (Figure S3A-C). This revealed no major changes in interaction partners, including the retention of tubulin binding, consistent with proficient meiotic spindle formation in *stu1ΔTOGL1* cells (Figure 2F and G). We note that Mad3^BUBR1^ was not recovered in either Stu1-GFP or Stu1ΔTOGL1-GFP immunoprecipitates (Figure S3A-C), indicating that only a minor fraction of cellular Stu1^CLASP^ interacts with Mad3^BUBR1^, while, conversely, the recovery of Stu1^CLASP^ in Mad3^BUBR1^ immunoprecipitates indicates that a major fraction of cellular Mad3^BUBR1^ is associated with Stu1^CLASP^. The failure of spindle formation in *pCLB2-STU1* cells precluded the segregation of homologs while *stu1ΔTOGL1* showed a modest impairment of homolog segregation in meiosis I, comparable to *mad3Δ* (Figure 2H). We conclude that *stu11TOGL11* is a separation of functional allele that loses Stu1^CLASP^ interaction with Mad3^BUBR1^, while retaining its ability to organise microtubules.

### Stu1^CLASP^ kinetochore localisation in meiosis does not require TOGL1

In mitosis, the TOGL1 domain recruits Stu1 to kinetochores where it is particularly enriched when microtubules are not attached^11^. However, we found in meiotic cells that Stu1ΔTOGL1-GFP was recruited to kinetochores similarly to wild-type Stu1-GFP (Figure S3D-I). Specifically, Stu1 was initially recruited to kinetochores at prophase exit before localizing to kinetochores and the spindle in metaphase I and II, or the spindle midzone in anaphase I and II (Figure S3D and E). Quantification of the kinetochore localization of Stu1ΔTOGL1-GFP compared to Stu1-GFP at prophase I exit, when kinetochores cluster prior to spindle formation, revealed a small, but not statistically significant, decrease (Figure S3F and G). Stu1ΔTOGL1-GFP also localized to unattached kinetochores in meiotic cells treated with the microtubule-depolymerising drug, benomyl (Figure S3H). Therefore, Stu1 association with kinetochores does not require its TOGL1 domain in meiosis. Consistently, Stu1 also localized to kinetochores independently of Mad3 (Figure S3I). Therefore, Stu1 is recruited to meiotic kinetochores independently of its TOGL1 domain or Mad3. The reason that the Stu1 TOGL1 domain is required in mitosis^11^ but not meiosis, is unclear but could reflect the distinct functional organisation of the meiotic kinetochore^13^.

### Mad3^BUBR1^ and the TOGL1 domain of Stu1^CLASP^ work together to promote meiosis I chromosome segregation

Our finding that *stu1ΔTOGL1* is a separation-of-function allele that specifically abolishes the Stu1^CLASP^-Mad3^BUBR1^ interaction provides a tool to address the function of this interaction. In particular, we aimed to test whether the checkpoint-independent function of Mad3^BUBR1^ in meiotic chromosome segregation is mediated through Stu1^CLASP^. If this is the case, we would predict that *mad31 stu11TOGL1* cells show similar meiosis I non-disjunction rates to either single mutant. We therefore live imaged cells with both copies of chromosome V labelled with tdTomato and carrying *GFP-TUB1*. As before, homologs disjoined to the same pole less than 5% of the time in *mad3Δ* cells, while disjunction was approximately doubled in *stu1ΔTOGL1* cells (Figure 3A). However, homolog mis-segregation in *mad31 stu11TOGL1* cells was similar to *stu1ΔTOGL1* alone (Figure 3A), suggesting that Mad3^BUBR1^ works in the same pathway as the TOGL1 domain of Stu1^CLASP^. Since Mad3^BUBR1^ function in meiosis I chromosome segregation is most evident in the absence of *MAD2* (Figure 1B), we assessed the *stu1ΔTOGL1* mutant in the *mad2Δ* background (Figure 3A). This revealed that *mad2Δ* exacerbated the meiosis I chromosome segregation defect in *mad3Δ* and *stu1ΔTOGL1* cells to a similar extent (Figure 3A). Importantly, however, we did not observe additional additive effects in the triple *mad21 mad31 stu11TOGL1* mutant. Therefore, Mad3^BUBR1^ and the TOGL1 domain of Stu1^CLASP^ act in the same genetic pathway to promote meiosis I homolog segregation. In contrast, Mad2 acts in a distinct pathway (Figure 1C and Figure 3A) and, unlike Mad3^BUBR1^, is important for homolog biorientation during meiosis I^4^ (see also below). Taken together, our findings indicate that Mad3^BUBR1^-Stu1^CLASP^ rescues the segregation of homologs that fail to biorient.

**Figure 3.**
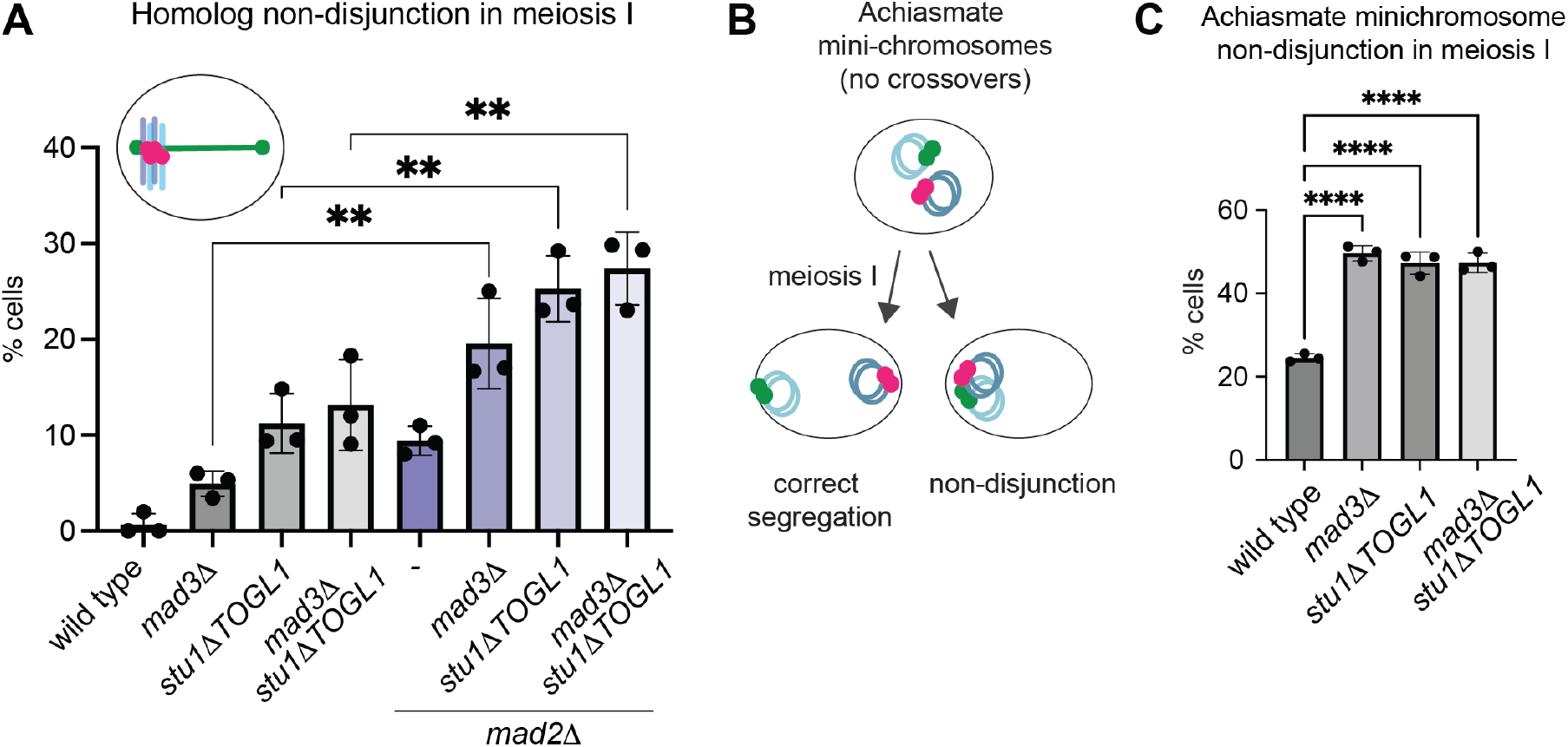
Mad3^BUBR1^ and the TOGL1 domain of STU1^CLASP^ rescue the segregation of chromosomes that fail to crossover or biorient. (A) Mad3-Stu1-TOGL1 and Mad2 act in distinct chromosome segregation pathways. Scoring of meiosis I non-disjunction after live imaging of endogenous homologous chromosomes competent for crossover recombination, as in Figure 1A-C. Mean of three biological replicates (wild type, *mad31, mad21* and *mad21 mad31* are as in 1C; *n=42-53 stu11TOGL1*; *n*=49-60 *mad31 stu11TOGL1*; *n*=47-65 *mad21 stu11TOGL1*; *n*=57-60 *mad21 mad31 stu11TOGL1*) where all genotypes were imaged concurrently is shown, with error bars representing standard deviation. ** p≤0.01, One way ANNOVA (Tukey’s multiple comparisons test). (B and C) Segregation of achiasmate minichromosomes requires Mad3 and the TOGL1 domain of Stu1. (B) Schematic showing segregation of GFP- (green) and tdTomato- (red) labelled achiasmate chromosomes in meiosis I. (C) Non-disjunction of achiasmate minichromosomes in the indicated genotypes. For each of three biological replicates, 50 anaphase I cells (as judged by DAPI staining) were scored after fixation. Bar chart shows mean with error bars representing standard deviation. **** p≤0.0001, One way ANNOVA (Tukey’s multiple comparisons test).

### The Mad3 and Stu1-TOGL1 segregation pathway rescues the segregation of chromosomes that lack crossovers

Crossover recombination generates chiasmata which provide the linkages that allow the generation of tension as homologs are pulled to opposite poles, ensuring accurate meiosis I segregation^3^. Accordingly, a failure to crossover is associated with severe aneuploidy and inviable gametes. However, in some circumstances, the absence of a crossover can be tolerated and such chromosomes which lack crossovers (also called achiasmate or non-exchange chromosomes) can undergo proper meiosis I segregation, though the underlying mechanisms are not well understood^14^. Even in systems where crossing over is typical, chromosomes that fail to crossover can segregate accurately. A single budding yeast achiasmate chromosome pair in a nucleus where all other chromosome pairs are linked by chiasmata segregates to opposite poles ~80% of the time^15–18^. Interestingly, Mad3^BUBR1^, unlike Mad2, is essential for the segregation of such an achiasmate chromosome pair^19^, though the underlying mechanism is unclear. Current models posit that the synaptonemal complex, which zips homologous chromosomes together as they recombine, persists at centromeres to maintain homolog pairing beyond prophase, thereby providing a backup mechanism that rescues the segregation of chromosomes that fail to crossover^20,21^. However Mad3^BUBR1^ is thought to act independently from centromere pairing, perhaps through mediating a prophase delay^19^. Our discovery that Mad3^BUBR1^ functions in a backup mechanism together with Stu1^CLASP^ to promote the segregation of normal chromosome pairs, which harbor chiasma, provided an alternative explanation. We hypothesised that Mad3^BUBR1^ directs achiasmate chromosome segregation via engaging Stu1^CLASP^. To test this idea, we introduced a pair of centromeric minichromosomes, one labelled with tdTomato (*tetO-*TetR-tdTomato), the other with GFP (*lacO-*GFP-LacI) into *mad3Δ, stu1ΔTOGL1* and *mad31 stu11TOGL1* cells (Figure 3B). Since the mini-chromosomes are both small and divergent in sequence, they will not crossover and therefore represent a pair of achiasmate chromosomes. In wild-type cells, red and green signal segregated to the same pole at anaphase I in 25% of cells, while segregation of the achiasmate mini-chromosomes was essentially random (~50%) in *mad3Δ* cells, as expected^19^ (Figure 3C). Crucially, *stu1ΔTOGL1* cells, where the Mad3^BUBR1^-Stu1^CLASP^ interaction is abolished (Figure 2B-D), also exhibit random segregation of achiasmate mini-chromosomes in meiosis I, as does the *mad31 stu11TOGL1* double mutant (Figure 3C). Therefore, the ability of Stu1^CLASP^ to bind Mad3^BUBR1^ is critical for achiasmate chromosome segregation.

### Mad3^BUBR1^ enables chromosome-spindle interactions through Stu1^CLASP^

How might Stu1^CLASP^-Mad3^BUBR1^ contribute to the fidelity of chromosome segregation? Stu1 and CLASP proteins are known to regulate microtubule dynamics at kinetochores and thereby promote stable kinetochore capture and biorientation^11,22–24^. Therefore, Mad3^BUBR1^ may promote kinetochore capture or biorientation via Stu1^CLASP^. To test this, we analysed the position of *CEN5-*tdTomato foci relative to the metaphase I spindle prior to elongation at anaphase I (Figure 4A-C). We considered instances where *CEN5*-tdTomato was located centrally on the spindle axis as “bioriented”, while asymmetric *CEN5*-tdTomato foci on the spindle axis were scored as “off centre” and foci which did not co-locate with the GFP-Tub1 signal were scored as “off axis” (Figure 4A). Consistent with a previous report^4^, *mad2Δ* showed defective biorientation, manifest as a significantly increased frequency of “off centre” *CEN5*-tdTomato foci (Figure 4B). However, neither *mad3Δ* nor *stu1ΔTOGL1* exhibited defective biorientation unless *MAD2* was also deleted (Figure 4B). In contrast, the fraction of cells where *CEN5*-tdTomato were “off axis”, suggesting defective kinetochore capture by microtubules or stabilisation of this attachment, was significantly increased over wild type in both *mad3Δ* and *stu1ΔTOGL1* mutants whether or not Mad2 was present (Figure 4C). These data show that Mad2 is important for positioning chromosomes in the centre of the spindle axis (bioriented), while Stu1^CLASP^-Mad3^BUBR1^ is important for chromosome association with the spindle. Therefore, Mad2 and Stu1^CLASP^-Mad3^BUBR1^ represent two distinct chromosome segregation pathways that respectively promote biorientation and chromosome-microtubule interactions, possibly through initial capture.

**Figure 4.**
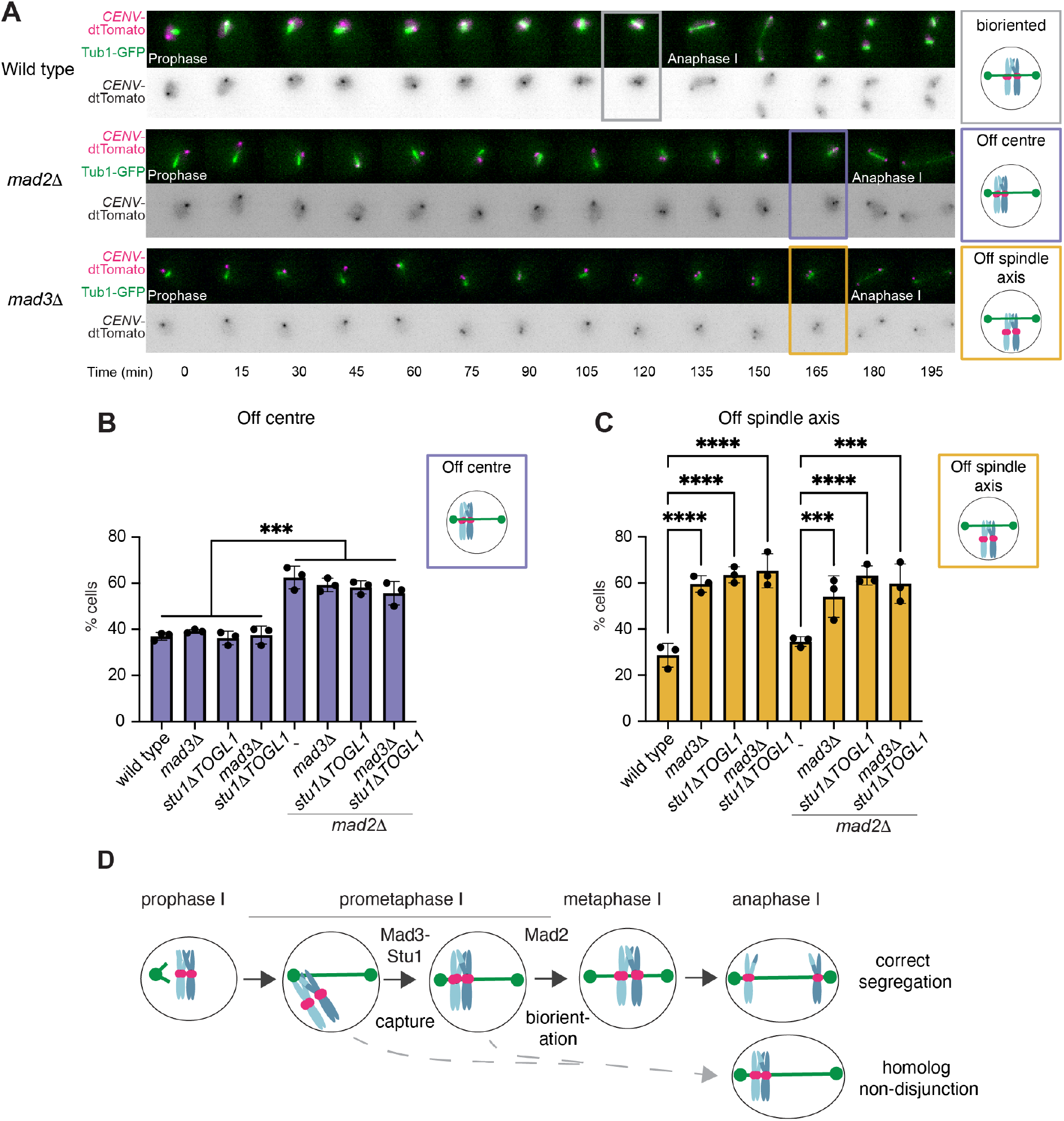
Mad3^BubR1^-Stu1^CLASP^ promote chromosome association with spindles during meiosis I. (A-C) Live imaging reveals position and orientation of *CEN5-GFP* relative to the metaphase I spindle (*GFP-TUB1*). (A) Representative time series showing chromosome capture and alignment on the meiosis I spindle for the indicated genotypes and scenarios. (B and C) The percentage of cells where *CEN5*-GFP foci (which typically were observed as one or two foci) were located off the centre of the spindle (B) or off the spindle axis (C) was scored in the last time point prior to anaphase I spindle elongation. Examples of correctly bioriented (grey box), off centre (purple box) or off axis (yellow box) are shown in A. Bar charts show mean of three biological replicates (*n*=44-56 wild type; *n*=52-62 *mad31*, *n*=36-52 *stu11TOGL*; *n*=41-49 *mad31 stu11TOGL1*; *n*=45-58 *mad21*; *n*=41-56 *mad21 mad31*; *n*=28-54 *mad21 stu11TOGL1*; *n*=25-58 *mad21 mad31 stu11TOGL1*) with error bars representing standard deviation. **** p<0.0001, ***p≤0.001, one way ANNOVA (Tukey’s multiple comparisons test). (D) Model for the steps leading to homolog alignment in meiosis I. First, Mad3 engages Stu1 to facilitate chromosome association with microtubules. Second, Mad2 aligns chromosomes in the centre of the spindle through an unknown mechanism.

## Conclusions

The spindle checkpoint has long been recognised as a critical surveillance mechanism that prevents catastrophic segregation where kinetochores are not properly attached to microtubules. Here we provide evidence that in addition to their checkpoint role, the components of this surveillance mechanism also contribute directly to the correction of improper or absent kinetochore-microtubule attachments. Incorporating these two key activities within an individual protein provides an opportunity to coordinate surveillance with segregation mechanisms. The existence of two separate segregation-promoting branches of the spindle checkpoint proteins also provides the cell with additional opportunities for coordination and safeguarding. While Mad2 promotes sister kinetochore biorientation, we identify a new pathway of chromosome alignment whereby Mad3^BUBR1^ engages the Stu1^CLASP^ protein in capturing kinetochores into the spindle. We demonstrate that this pathway is critical to rescue chromosomes from lack of crossovers or a failure to biorient. Meiotic errors contributing to aneuploidy in human oocytes are predicted to originate from multiple deficiencies that accumulate as women age. Safeguarding mechanisms such as Mad3^BUBR1^-Stu1^CLASP^-dependent kinetochore capture are therefore likely to play key roles in protecting against such errors.

## Acknowledgements

We are grateful to Kevin Hardwick for helpful discussions, to Alexander Julner Dunn, Lori Koch, Lucia Massari and Gerard Pieper for comments on the manuscript. We thank Tomo Tanaka, Wolfgang Zachariae and Eris Duro for plasmids and yeast strains. We gratefully acknowledge the Wellcome Centre Optical Imaging Laboratory (COIL) and Proteomics Facility for microscopy and mass spectrometry support, respectively. This work was funded through a Wellcome Investigator award to ALM [220780] and core funding for the Wellcome Centre for Cell Biology [203149]. AM also gratefully acknowledges studentship support from the Darwin Trust of Edinburgh.

## Author contributions

Conceptualization: ALM and AM; Methodology: AM & CS; Formal analysis: AM & CS; Investigation: AM; Writing – Original draft: ALM; Writing – Review & Editing: AM, CS & ALM. Visualization: AM and ALM; Supervision: ALM; Funding acquisition: ALM.

## Methods

### Materials, data and code availability

Further information and requests for resources and reagents should be directed to and will be fulfilled by the lead contact, Adele Marston. Yeast strains and plasmids used in this study can be obtained from the lead contact, without restriction. Mass spectrometry datasets reported in this study have been deposited at PRIDE and are publicly available as of the data of publication. DOIs are in the key resources table.

The paper does not report original code.

Any additional information required to reanalyze the data reported in this paper is available from the lead contact upon request.

### Yeast strains

All yeast strains are SK1 derivatives and are listed in Table S1. Gene deletions, promoter replacements and gene tags were introduced using standard PCR-based methods^26^. *pCLB2-CDC20* ^27^, inducible-*ndt80 (pGAL1-NDT80*, *pGPD1-GAL4.ER*; ^28^), *CEN5*-GFP^29^ *CEN5-tdTomato*^30^ and *GFP-TUB1*^30^ were described previously and *stu1ΔTOGL1* was made by CRISPR-Cas9 in this study.

### Plasmids

For the achiasmate minichromosome assay we used plasmid p331^31^ (*CEN4, URA3, tetO* arrays) and plasmid AMp1963. Plasmid AMp1963 was generated in this study by restriction digest of yCPlac22 and pAFS59 with *Sal*I and *Kpn*I and ligating the respective 10,108bp *lacO* and 4,743 bp *CEN4 TRP1* fragments.

### Growth conditions

#### Meiotic induction and growth of yeast cultures

Diploids were recovered from 20% glycerol stock at −80°C onto YPG plates (1% yeast extract, 2% bactopeptone, 2% glycerol, and 2% agar) and grown for 12h. They were patched on 4% YPDA agar plates (1% yeast extract, 2% bactopeptone, 4% glucose, and 2% agar) for 6-8h, then inoculated in YPDA medium (1% yeast extract, 2% bactopeptone, 2% glucose, 0.3 mM adenine) and shaken at 250rpm at 30°C for approximately 24h before being diluted to OD_600_ = 0.2-0.4 in YPA (1% yeast extract, 2% peptone, 1% potassium acetate) and grown for another 12-16h. In the morning, cells were pelleted and washed with sterile dH_2_O before being resuspended in SPO media (0.3% potassium acetate) at OD_600_ = 1.8-1.9 and shaken at 250rpm at 30°C for the duration of the experiment. Cells grown for live-cell imaging were resuspended at OD_600_ = 2.3 and for mass spectrometry experiments at OD_600_ 2.5 in SPO media. For synchronous meiosis, inducible-*NDT80* was used to allow prophase I block-release^32^. For prophase I arrest, cells with inducible-*NDT80* were harvested after 6h in SPO medium. For metaphase I arrest, cells with *pCLB2-CDC20* were harvested after 6h in SPO medium.

#### Sporulation efficiency assay

Diploids were recovered on YPG plates overnight and sporulated in liquid SPO media at 250rpm at 30°C for 72h and 200 cells were scored by light microscopy to determine the proportion of triads and tetrads, dyads and non-sporulated cells.

### Mass spectrometry methods

#### Conjugating anti-FLAG or anti-GFP to dynabeads

Protein G Dynabeads (Invitrogen) were washed twice in 1ml 0.1M Na-phosphate, pH 7.0, before incubating with 1/10^th^ volume of M2 anti-FLAG monoclonal antibody (Sigma) or 1/5^th^ volume of anti-GFP antibody (Roche) and 50μl of 0.1M Na-phosphate with gentle agitation for 30min at room temperature. Beads were washed twice in 1ml of 0.1M Na-phosphate pH 7.0 with 0.01% Tween-20, then washed twice with 1ml of 0.2 M triethanolamine, pH 8.2. Antibody-conjugated Dynabeads were resuspended in 1ml of 20mM DMP (Dimethyl Pimelimidate, D8388, Sigma) in 0.2M triethanolamine, pH8.2 (prepared immediately before use) and incubated with rotational mixing for 30min at room temperature. Beads were concentrated, the supernatant removed and 1ml of 50mM Tris-HCl, pH7.5 added before incubating for 15min with rotational mixing. The supernatant was removed and beads were washed three times with 1ml 1XPBST+0.1% Tween-20 before resuspending in 300ml of 1xPBST.

#### Immunoprecipitation

Either 3L (Figure 1E-H) or 200ml (Figures 2C-D, S2B, and S3A-C) of meiotic culture grown at OD_600_= 2.5 was harvested and washed once with sterile dH_2_O. Cells were pelleted and resuspended in 20% v/w 2 mM PMSF and snap frozen as small ‘noodles’ by releasing drops of cells into liquid nitrogen. These noodles were filled in metal canisters pre-cooled in liquid nitrogen and cells lysed by 5 rounds of 30/s speed for 3 min each in the twin bio-pulveriser Retsch MM400. Grindate was then emptied out of the canisters into a 50ml falcon tube and stored in −80°C. For immunoprecipitation, the cryogrindate was thawed and resuspended in 20% w/v H0.15M lysis buffer (25mM Hepes pH8, 2mM MgCl_2_, 0.1mM EDTA pH8.0, 0.5mM EGTA-KOH pH8.0, 15% glycerol, 0.1% NP-40, 150mM KCl) with phosphatase and protease inhibitors (CLAAPE, comprising 10 μg/ml each of **<u>c</u>**hymostatin, **l**eupeptin, **<u>a</u>**ntipain, **<u>p</u>**epstatin and **<u>E</u>**64, together with 2mM AEBSF, 0.8mM Na Orthovanadate, 0.2uM microcystin, 1x EDTA-free Roche protease inhibitor tablet, 2mM NEM, 4mM β-glycerophosphate, 2mM Na pyrophosphate, 10mM NAF). 40U/ml of Benzonase (Novagen) was added to the lysate and incubated for 1h at 4°C with rotation to digest DNA. Samples were centrifuged for 10min at 4000rpm at 4°C and supernatant was collected in new pre-chilled falcon tubes. Protein concentration was determined by Bradford assay, and each lysate was adjusted to the same volume and protein concentration. 50μl of each adjusted lysate was added to 10μl 4xLDS + 5% β-mercaptoethanol, boiled at 95°C for 5min and stored at −20°C as input. 2μg ⍺-GFP (Roche) or 0.05 μg ⍺-FLAG (Sigma) previously conjugated to Protein G-dynabeads were added to each sample and incubated with rotation at 4°C for 2.5h. Dynabeads were concentrated using a pre-chilled magnet and the flow through was discarded. The beads were transferred an eppendorf tube and washed once with buffer H0.15M with inhibitors and 2mM DTT, then three more times with buffer H0.15M with inhibitors. Beads were concentrated on the magnet, resuspended in 50μl 1xLDS + 5% β-mercaptoethanol and boiled at 70°C for 10min to elute. Samples were spun down at 13,200rpm for 5min before the eluate was transferred to a fresh eppendorf tube and stored at −20°C indefinitely for preparation for mass spectrometry.

#### In gel digestion of protein samples for mass spectrometry

In-gel digestion was used to prepare samples for mass spectrometry in Figure 1E-H and 2B. Yeast growth conditions and the immunoprecipitation protocol used was the same as above, with a few modifications. 3L of SPO cultures at OD_600_=2.5 were harvested and 500μl Protein G dynabeads previously conjugated to 50μl M2 ⍺-FLAG antibody were added to each extract which was made from approximately 15g of cryogrindate. Proteins were eluted from beads in 100μl 1xLDS + 5% β-mercaptoethanol, out of which 90μl was loaded on NuPAGE Novex 4-12% Bis-Tris Gel (Life Technologies) gels and run for 6min so that all proteins enter the gel. The gel was stained by incubating with agitation in Instant Blue (Abcam) and washed three times for 5min each with dH_2_O. Protein bands were cut from the gel and chopped into ~1mm^3^ pieces using a new clean scalpel, and the pieces were collected in an eppendorf tube. The pieces were submerged in 50mM ammonium bicarbonate (ABC) for 30min. ABC was discarded and 100% acetonitrile (ACN) was added until gel pieces were submerged and incubated for another 30min. ~80ul 10mM DTT in 50mM ABC was added to the gel pieces and incubated for 30min at 37°C. DTT solution was removed and gel pieces were resuspended in ACN for 5min and any excess liquid was removed. ~80ul 55mM iodoacetamide dissolved in ABC was added to cover the pieces and incubated in the dark at RT for 20mins. The liquid was removed and gel pieces were incubated with 50mM ABC buffer for 5min at 37°C, the ABC was removed, and then the gel pieces were incubated in ACN for 5mins at 37°C. All liquid was removed and enough trypsin digestion mix (0.013μg/ml trypsin, 10% ACN, 10mM ABC) was added to cover the gel pieces and left initially at 4°C and then at 37°C overnight for 12-15h in a moist chamber. 0.1% or 10% TFA was added to the gel pieces in trypsin digestion mix to stop over-digestion of peptides and the solution was kept at room temperature for 15min to allow all peptides to diffuse out form the gel. 1μl of sample was dropped on a pH paper to confirm that the solution has pH <2.0.

#### Filter-aided Sample Preparation (FASP) of protein samples

FASP was used to prepare samples for mass spectrometry in Figure 2C-D, S2B, and S3A-C, as described^33^, with a few modifications. Proteins were eluted from beads by incubating in 30μl 0.1% Rapigest (Waters) dissolved in 50mM ABC at 37°C for 30min, removing the eluate and then repeating to obtain a further 30μl of eluate. Pooled eluates from the two elutions were stored at −20°C. On the day of trypsin digestion, 10% volume of 1M DTT was added to samples and boiled for 5min at 95°C with agitation. Tubes were cooled to room temperature before adding 3x vol of 8M urea in 100mM Tris-HCl pH8.0 (UBB) to each sample. The whole sample was transferred onto Sartorius Stedim Biotech’s Vivacon 500 MWCO 30 000 VN01H21 column and spun down at 10,000rpm for 10-15min at room temperature to bind all peptides to the membrane. 100μl of 55mM iodoacetamide dissolved in UBB was added, the tube shaken at 600rpm for 1min at RT in a theromixer, and incubated in the dark for 30min before spinning the buffer through the column. The column was then washed once with 100μl UBB and twice with 100μl ABC. The column was completely dried before adding 60μl of trypsin digestion mix (0.013μg/ml trypsin, 0.002% TFA, 50mM ABC) onto the column membrane. Columns were capped and sealed with parafilm before being shaken at 600rpm for 1min at room temperature and then incubated at 37°C overnight for ~15h in a moist chamber. Parafilm was removed and the columns were centrifuged to elute trypsin-digested peptides into new protein protein LoBind tubes containing 10μl of 10% TFA to stop the trypsin digestion. 1μl of sample was dropped on a pH paper to confirm that the solution was pH <2.0.

#### Mass spectrometry

Stage tips were prepared by inserting three Empire C18 disks (3M) inside a p200 pipette tip. 20μl MeOH and 50μl 0.1% TFA was passed through the tip to calibrate the disks at the correct pH. All the liquid from the gel digestion or the in-column digestion was passed through the stage tip by microfuging for ~10min. The tip was then washed again with 0.1% TFA and stored in −20°C. Peptides were eluted in 40 μL of 80% acetonitrile in 0.1% TFA and concentrated down to 1 μL by vacuum centrifugation (Concentrator 5301, Eppendorf, UK). The peptide sample was then prepared for LC-MS/MS analysis by diluting it to 6 μL by 0.1% TFA.

All LC-MS analyses were performed on an Orbitrap Fusion™ Lumos™ Tribrid™ Mass Spectrometer (Thermo Fisher Scientific, UK) both coupled on-line, to an Ultimate 3000 HPLC (Dionex, Thermo Fisher Scientific, UK). Peptides were separated on a 50 cm (2 µm particle size) EASY-Spray column (Thermo Scientific, UK), which was assembled on an EASY-Spray source (Thermo Scientific, UK) and operated constantly at 50°C. Mobile phase A consisted of 0.1% formic acid in LC-MS grade water and mobile phase B consisted of 80% acetonitrile and 0.1% formic acid. Peptides were loaded onto the column at a flow rate of 0.3 μL min^-1^ and eluted at a flow rate of 0.25 μL min^-1^ according to the following gradient: 2 to 40% mobile phase B in 150 min and then to 95% in 11 min. Mobile phase B was retained at 95% for 5 min and returned back to 2% a minute after until the end of the run (190 min).

Survey scans were recorded at 120,000 resolution (scan range 350-1500 m/z) with an ion target of 4.0e5, and injection time of 50ms. MS2 was performed in the ion trap at a rapid scan mode, with ion target of 2.0E4 and HCD fragmentation^34^ with normalized collision energy of 27. The isolation window in the quadrupole was 1.4 Thomson. Only ions with charge between 2 and 6 were selected for MS2. Dynamic exclusion was set at 60s.

#### Analysis of mass spectrometry data

The MaxQuant software platform^35^ version 1.6.1.0 was used to process the raw files and search was conducted against the complete/reference proteome set of *Saccharomyces cerevisiae* SK1 strain (combined Saccharomyces Genome Database and in-house database - released in August 2019), using the Andromeda search engine^36^. For the first search, peptide tolerance was set to 20 ppm while for the main search peptide tolerance was set to 4.5 pm. Isotope mass tolerance was 2 ppm and maximum charge to 7. Digestion mode was set to specific with trypsin allowing maximum of two missed cleavages. Carbamidomethylation of cysteine was set as fixed modification and oxidation of methionine, was set as variable modification. Label-free quantitation analysis was performed by employing the MaxLFQ algorithm as described^37^. Absolute protein quantification was performed as described^38^. Peptide and protein identifications were filtered to 1% FDR.

Statistics from LFQ data were processed using Bioconductor *DEP* R package according to^39^(https://github.com/arnesmits/DEP).

### Microscopy methods

#### Live cell imaging

To adhere cells, 5μl of ConA (5mg/mL ConcanavalinA in 50mM CaCl_2_, 50mM MnCl_2_) was spread at the bottom of chambers in 8-well glass-bottomed Ibidi dish (Thistle Scientific) using a plastic loop and incubated in 30°C for 15min. ConA was aspirated and the chambers washed three times with 500μl sterile dH_2_O and stored in the dark. To prepare cells for imaging, 10ml of meiotic cultures were started at OD_600_=2.3 in SPO media. After 3h, 1ml of culture was spun down at 3000rpm for 1min. The pellet was resuspended in 300μl SPO media, added to the Ibidi dish and incubated for 20min at 30°C. Wells were washed with 500μl SPO media twice before adding 400μl fresh SPO media. For the inducible Ndt80 block-release system, 200μl SPO was added while setting up the Ibidi dish, and another 200μl SPO with 2μM β-estradiol was added immediately before starting the time lapse imaging. For depolymerizing microtubules in Figure S3H, benomyl was pre-dissolved in SPO medium and was added at a final concentration of 50μg/ml to the wells along with β-estradiol. Fluorescent microscopy was performed using Zeis Axioplan 2 microscope with 100x Plan ApoChromat NA 1.4 oil lens. Images were acquired through ORCA FLASH 4 CCD camera with auto-focus operated through Axiovision software and with 2×2 binning. GFP-Tub1 was imaged at 4% laser intensity for 80ms, *CEN5*-tdTomato was imaged at 4% intensity for 100ms, Mtw1-tdTomato was imaged at 10% intensity for 100ms. For all fluorescent channels, 9 z-slices of 0.7μm interval were captured. Brightfield was used for auto-focus and imaged only for the middle slice with 3V for 10ms. Chromosome segregation assays were imaged every 15min and biorientation assays were imaged every 5min for 10h in total.

#### Image analysis

ImageJ software (National Institutes of Health) was used to max project the z-stacks and for visualising the images. To quantify signal intensity in Figure S3G, a circular region was drawn encompassing the region of GFP and tdTomato signal overlap, and the ratio of integrated density measurement of the GFP signal over the tdTomato signal was calculated. Final image assembly was conducted in Adobe Illustrator.

#### Imaging of GFP-labelled chromosomes in fixed cells

For the chromosome segregation assay reported in Figure S1, 150μl meiotic culture at OD_600_=1.9 in SPO media was added to 15μl of 37% v/v formaldehyde in 1.5ml Eppendorf tubes and fixed for 8min at room temperature. Tubes were then spun at 13,200rpm for 1min, supernatant removed and resuspended in 1ml 80% EtOH. Tubes were spun again for 30s, EtOH poured out, spun again for 15s and the remaining EtOH removed with a pipette. The pellet was resuspended in 20μl of 1μg/ml DAPI and temporarily stored at 4°C for up to one week. 3μl of cells were placed on a Superfrost microscope slide (Thermo Fisher Scientific), covered with a coverslip (VWR) and sealed with nail polish. The coverslip was pressed tightly against the slides and imaged on an Axioplan 2 microscope with 100x Plan ApoChromat NA 1.4 oil lens with 5% GFP, 10% tdTomato and 2% DAPI to visualise GFP and tdTomato dots and DNA.

#### Achiasmate minichromosome segregation assay

Diploid cells carried homozygous *pURA3-GFP-LacI* and heterozygous *pURA3-TetR-tdTomato* integrated into the genome, and AMp1963 and AMp1973 plasmids. The plasmids were maintained by selecting on synthetic complete glucose agar lacking both uracil and tryptophan (SD/-ura/-trp), before cells were inoculated consecutively in YPDA, YPA and SPO liquid media was described above for induction of meiosis and sporulation. Samples were collected 2h after inducing sporulation and every 30min thereafter until 4:30h. Cells were fixed and processed as described abov<u>e “Imaging of GFP-labelled chromosomes in fixed cells”,</u> <u>stored in 4°C</u> and were counted on the same day. Only cells with a single GFP and single tdTomato focus at the binucleate stage were included in segregation scoring.

#### Immunofluorescence

200μl of meiotic culture was centrifuged at 13,200rpm for 1min and resuspended in 500μl of 3.7% v/v formaldehyde in 0.1M KPi, pH6.4 (potassium phosphate buffer: 27.8mM K_2_HPO_4_ and 72.2mM KH_2_PO_4_) to fix overnight at 4°C. Cells were then spun down and washed three times with 1ml of 0.1M KPi buffer, and resuspended in 1ml of sorbitol-citrate (1.2M sorbitol, 0.1M KH_2_PO_4_, 36mM citric acid). Cells were spun down again and resuspended in digestion mix (200μl sorbitol-citrate, 20μl glusulase and 6μl 10mg/ml zymolase) and incubated at 30°C for 2h or until cells become phase-dark under light microscope. Once digested, cells were pelleted, washed with 1ml sorbitol-citrate and then resuspended in ~50μl sorbitol-citrate. 5μl of 0.1% polylysine was added to each well of multi-well slides (Thermo Fisher Scientific) for 5min at room temperature before being rinsed with dH_2_O and air-dried. 5μl of digested cells were added to each well and incubated for 10min, before aspiration and submerging in MeOH for 3min followed by acetone for 10s. 5μl of Rat ⍺-tubulin (AbD Serotec) primary antibody diluted in 1:50 in PBS-BSA (1% w/v BSA, 0.04M K_2_HPO_4_, 0.01M KH_2_PO_4_, 0.15M NaCl, 0.1% w/v NaN_3_) was added to each well and incubated in a moist chamber for 1h at room temperature. Primary antibody was aspirated and wells washed five times each with 5μl of PBS-BSA. 5μl of Donkey anti-rat-FITC (Jackson ImmunoResearch) secondary antibody was added, and slides were incubated in a dark moist chamber for 1h, then each well was washed five times with 5μl of PBS-BSA. 3μl of DAPI-mount (9mM p-phenylenediamine, 0.04M K_2_HPO_4_, 0.01M KH_2_PO_4_, 0.15M NaCl, 0.1% w/v NaN_3_, 50ng/ml DAPI, 90% v/v glycerol) was added to each well, before the slide was covered with a glass coverslip and sealed with nail paint. Slides were stored at −20°C and visualized on a Zeiss Axioplan 2 microscope with 100x Plan ApoChromat NA 1.4 oil lens.

## Quantification and statistical analysis

Statistical analysis and graphs were generated using Graphpad Prism 9 software (San Diego). Micrographs and graphs were assembled using Adobe Illustrator. Statistical details of all experiments are given in the figure legends.

## Supplemental information

**Figure S1.**
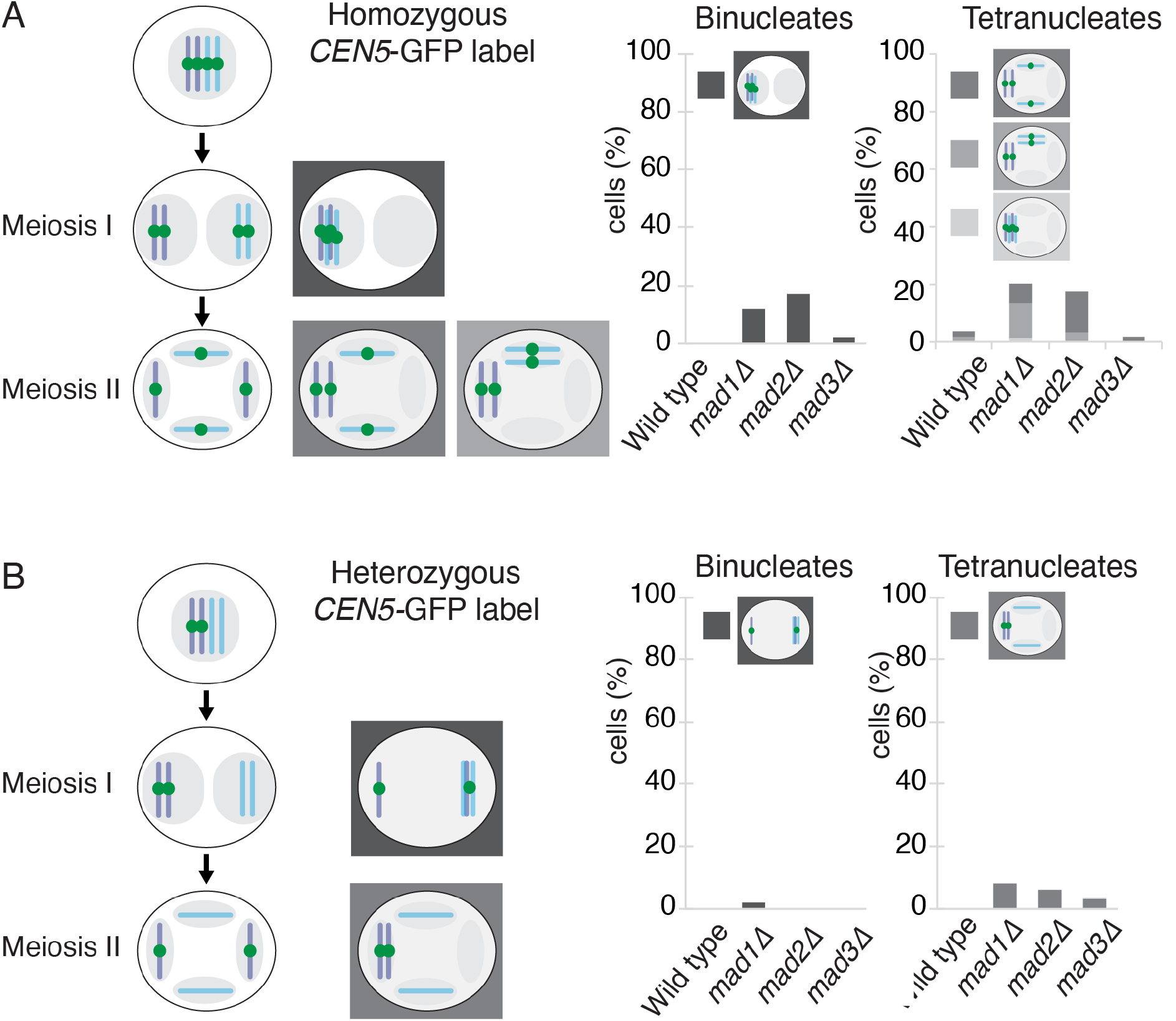
Mad2 is more critical than Mad3 for homolog segregation. **Related to Figure 1.** (A) Analysis of meiosis I and II segregation in *mad1Δ, mad2Δ* and *mad3Δ* cells carrying GFP label on both copies of chromosome V. Strains carrying homozygous *CEN5-GFP* were induced to sporulate, fixed at hourly intervals and counter-stained with DAPI. The pattern of chromosome segregation as shown in the schematic (left) was scored in 100 binucleate and 100 tetranucleate cells (right). (B) Analysis of meiosis I and II segregation in *mad1Δ, mad2Δ* and *mad3Δ* cells carrying GFP label on one copy of chromosome V. Strains carrying heterozygous *CEN5-GFP* were induced to sporulate and fixed at hourly intervals. The pattern of chromosome segregation as shown in the schematic (left) was scored in 100 binucleate and 100 tetranucleate cells (right).

**Figure S2.**
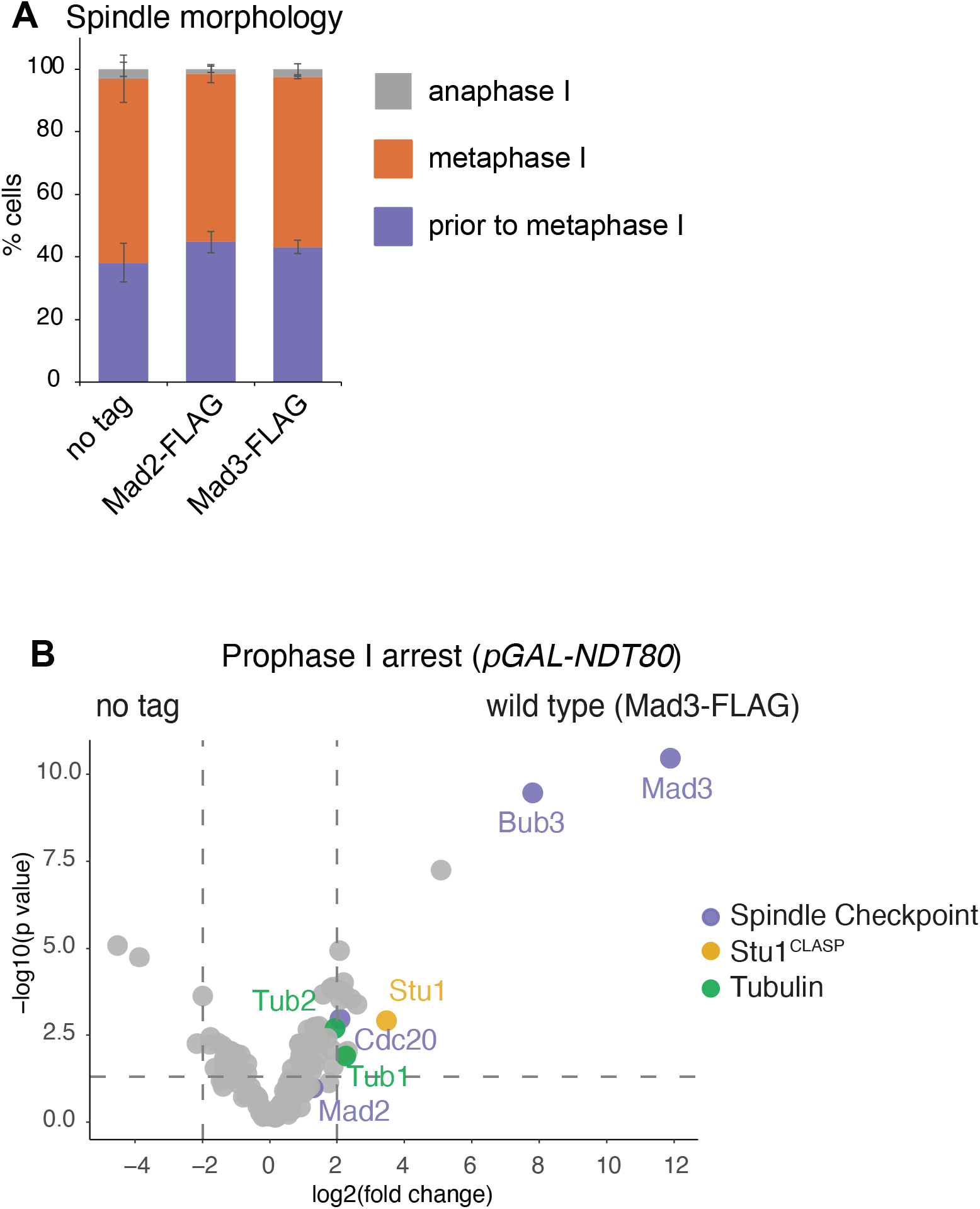
Mad2 and Mad3 interacting proteins in prometaphase I of meiosis. **Related to Figure 1.** (A) Confirmation that cells harvested for Mad2-FLAG and Mad3-FLAG shown in Figure 1E-G were in prometaphase/metaphase I. Spindle morphology after anti-tubulin immunofluorescence was scored in 200 cells of each replicate. Mean of three biological replicates with error bars indicating standard deviation. (B) Stu1^CLASP^ also associates with Mad3 in meiotic prophase I (*pGAL-NDT80* arrest). Volcano plots after mass spectrometry showing the relative enrichment of proteins immunoprecipitated with Mad3-FLAG in wild-type vs no tag. Data shown is from the same experiment as shown in Figure 2C and D.

**Figure S3.**
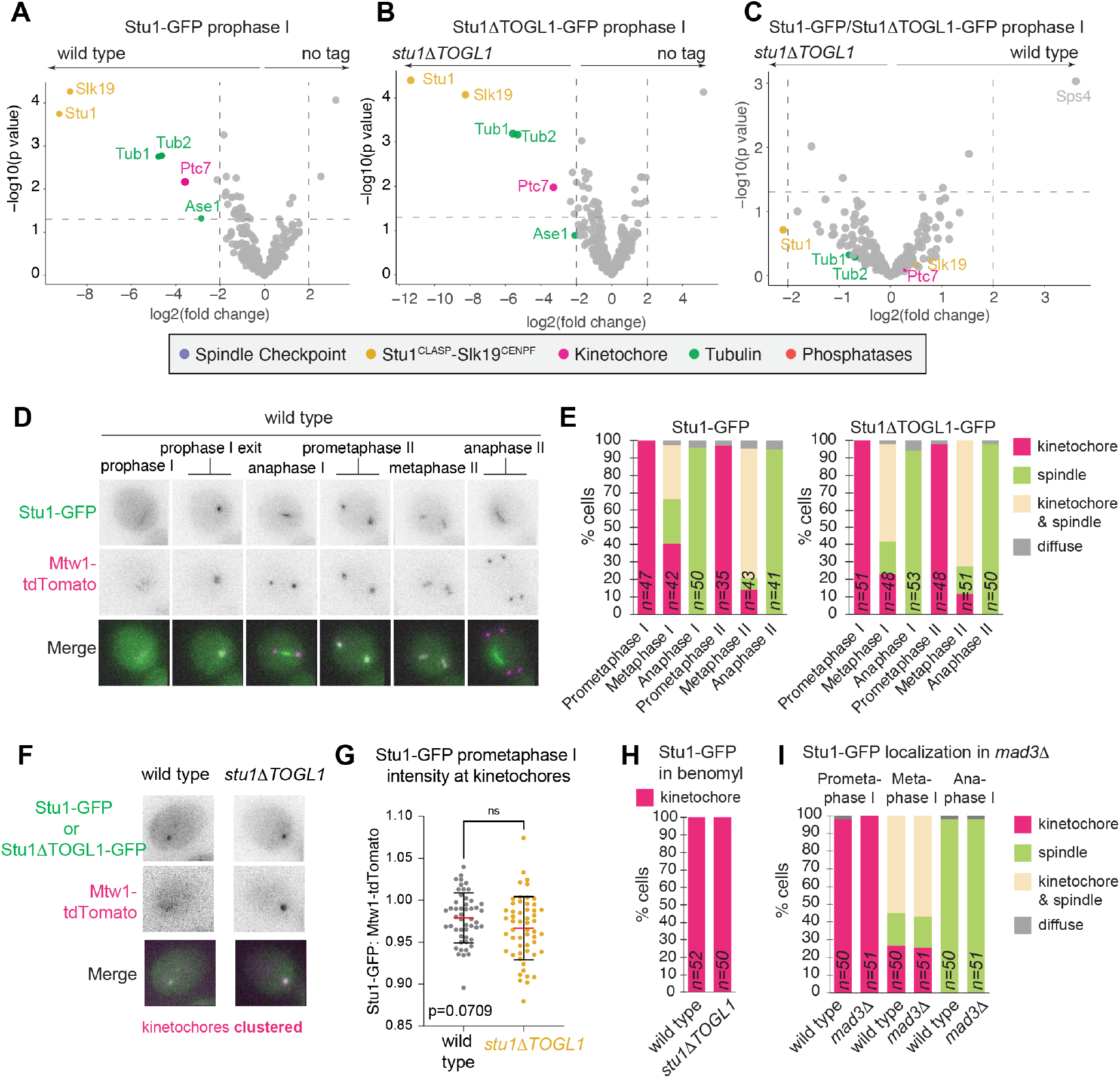
The Stu1 TOGL1 domain is not required for association with tubulin, Slk19 or kinetochores. (A-C) Tubulin and Slk19 are highly enriched in Stu1 immunoprecipitates independently of the TOGL1 domain. Stu1-GFP or Stu1ΔTOGL1-GFP anti-GFP immunoprecipitation followed by mass spectrometry was performed from cells harvested in a prophase I arrest (*GAL-NDT80*) Volcano plots show relative enrichments from three biological replicates and Log_2_(Fold Change) between conditions is shown with corresponding p values. Dashed line indicates Log_2_(Fold Change) = |2|. (A) Wild type Stu1-GFP vs no tag, (B) Stu1ΔTOGL1-GFP vs no tag and (C) Stu1-GFP vs Stu1ΔTOGL1-GFP. (D and E) Stu1 localization is unaffected in the absence of its TOGL1 domain. (D-G) Stu1 kinetochore and spindle localization does not require the TOGL1 domain. Cells where *STU1* is replaced by either Stu1-GFP (wild type) or Stu1ΔTOGL1-GFP (*stu1ΔTOGL1*) were induced to enter meiosis and live imaged in the presence of DMSO (D-G) or benomyl (H). (D) Representative images of Stu1-GFP in the indicated meiotic stages. (E) Quantification of Stu1 localization at the indicated stages. (F and G) The ratio of intensity of GFP vs tdTomato signal was determined upon kinetochore clustering at prophase exit. (F) Representative images. (G) Stu1ΔTOGL1-GFP intensity is not significantly reduced at kinetochores compared to Stu1-GFP intensity. Red line indicates mean intensity, bars represent 95% CI. 50 cells were measured for each condition. p=0.709 Mann-Whitney test (I) Mad3 is not required for Stu1 localization to kinetochores or spindles. Live cell imaging of wild type and *mad3*1 cells undergoing meiosis was quantified as in D and E.

## References

1. Duro, E., and Marston, A.L.A.L. (2015). From equator to pole: splitting chromosomes in mitosis and meiosis. Genes & development 29, 109–122. 10.1101/gad.255554.114.

2. McAinsh, A.D., and Kops, G.J.P.L. (2023). Principles and dynamics of spindle assembly checkpoint signalling. Nat Rev Mol Cell Biol, 1–17. 10.1038/s41580-023-00593-z.

3. Shonn, M.A., McCarroll, R., and Murray, A.W. (2000). Requirement of the spindle checkpoint for proper chromosome segregation in budding yeast meiosis. Science (New York, N.Y.) 289, 300–303.

4. Shonn, M.A., Murray, A.L.A.W., and Murray, A.L.A.W. (2003). Spindle checkpoint component Mad2 contributes to biorientation of homologous chromosomes. Current biology : CB 13, 1979–1984.

5. London, N., and Biggins, S. (2014). Mad1 kinetochore recruitment by Mps1-mediated phosphorylation of Bub1 signals the spindle checkpoint. Genes & development 28, 140–152. 10.1101/GAD.233700.113.

6. Sironi, L., Mapelli, M., Knapp, S., De Antoni, A., Jeang, K.-T., and Musacchio, A. (2002). Crystal structure of the tetrameric Mad1-Mad2 core complex: implications of a “safety belt” binding mechanism for the spindle checkpoint. EMBO J 21, 2496–2506. 10.1093/emboj/21.10.2496.

7. Luo, X., Tang, Z., Rizo, J., and Yu, H. (2002). The Mad2 Spindle Checkpoint Protein Undergoes Similar Major Conformational Changes Upon Binding to Either Mad1 or Cdc20. Molecular Cell 9, 59–71. 10.1016/S1097-2765(01)00435-X.

8. Hardwick, K.G., Johnston, R.C., Smith, D.L., and Murray, A.W. (2000). MAD3 encodes a novel component of the spindle checkpoint which interacts with Bub3p, Cdc20p, and Mad2p. The Journal of cell biology 148, 871–882.

9. London, N., Ceto, S., Ranish, J.A., and Biggins, S. (2012). Phosphoregulation of Spc105 by Mps1 and PP1 regulates Bub1 localization to kinetochores. Current biology : CB 22, 900–906.

10. Lawrence, E.J., Zanic, M., and Rice, L.M. (2020). CLASPs at a glance. J Cell Sci 133, jcs243097. 10.1242/jcs.243097.

11. Funk, C., Schmeiser, V., Ortiz, J., and Lechner, J. (2014). A TOGL domain specifically targets yeast CLASP to kinetochores to stabilize kinetochore microtubules. J Cell Biol 205, 555–571. 10.1083/jcb.201310018.

12. Pasqualone, D., and Huffaker, T.C. (1994). STU1, a suppressor of a beta-tubulin mutation, encodes a novel and essential component of the yeast mitotic spindle. J Cell Biol 127, 1973–1984. 10.1083/jcb.127.6.1973.

13. Borek, W.E., Vincenten, N., Duro, E., Makrantoni, V., Spanos, C., Sarangapani, K.K., de Lima Alves, F., Kelly, D.A., Asbury, C.L., Rappsilber, J., et al. (2021). The Proteomic Landscape of Centromeric Chromatin Reveals an Essential Role for the Ctf19 CCAN Complex in Meiotic Kinetochore Assembly. Current biology : CB 31, 283–296.e7. 10.1016/J.CUB.2020.10.025.

14. Kurdzo, E.L., and Dawson, D.S. (2015). Centromere pairing - Tethering partner chromosomes in meiosis i. FEBS Journal 282, 2445–2457. 10.1111/febs.13280.

15. Kemp, B., Boumil, R.M., Stewart, M.N., and Dawson, D.S. (2004). A role for centromere pairing in meiotic chromosome segregation. Genes & development 18, 1946–1951.

16. Dawson, D.S., Murray, A.W., and Szostak, J.W. (1986). An alternative pathway for meiotic chromosome segregation in yeast. Science (New York, N.Y.) 234, 713–717.

17. Mann, C., and Davis, R.W. (1986). Meiotic disjunction of circular minichromosomes in yeast does not require DNA homology. Proceedings of the National Academy of Sciences 83, 6017–6019. 10.1073/pnas.83.16.6017.

18. Guacci, V., and Kaback, D.B. (1991). Distributive disjunction of authentic chromosomes in Saccharomyces cerevisiae. Genetics 127, 475–488. 10.1093/genetics/127.3.475.

19. Cheslock, P.S., Kemp, B.J., Boumil, R.M., and Dawson, D.S. (2005). The roles of MAD1, MAD2 and MAD3 in meiotic progression and the segregation of nonexchange chromosomes. Nat Genet 37, 756–760.

20. Newnham, L., Jordan, P., Rockmill, B., Roeder, G.S., and Hoffmann, E. (2010). The synaptonemal complex protein, Zip1, promotes the segregation of nonexchange chromosomes at meiosis I. Proc Natl Acad Sci U S A 107, 781–785.

21. Gladstone, M.N., Obeso, D., Chuong, H., and Dawson, D.S. (2009). The synaptonemal complex protein Zip1 promotes bi-orientation of centromeres at meiosis I. PLoS genetics 5, e1000771.

22. Maiato, H., Sampaio, P., Lemos, C.L., Findlay, J., Carmena, M., Earnshaw, W.C., and Sunkel, C.E. (2002). MAST/Orbit has a role in microtubule–kinetochore attachment and is essential for chromosome alignment and maintenance of spindle bipolarity. Journal of Cell Biology 157, 749–760. 10.1083/jcb.200201101.

23. Maiato, H., Fairley, E.A.L., Rieder, C.L., Swedlow, J.R., Sunkel, C.E., and Earnshaw, W.C. (2003). Human CLASP1 Is an Outer Kinetochore Component that Regulates Spindle Microtubule Dynamics. Cell 113, 891–904. 10.1016/S0092-8674(03)00465-3.

24. Cheeseman, I.M., MacLeod, I., Yates, J.R. 3rd, Oegema, K., and Desai, A. (2005). The CENP-F-like proteins HCP-1 and HCP-2 target CLASP to kinetochores to mediate chromosome segregation. Current biology : CB 15, 771–777.

25. Perez-Riverol, Y., Csordas, A., Bai, J., Bernal-Llinares, M., Hewapathirana, S., Kundu, D.J., Inuganti, A., Griss, J., Mayer, G., Eisenacher, M., et al. (2019). The PRIDE database and related tools and resources in 2019: Improving support for quantification data. Nucleic Acids Research 47, D442–D450. 10.1093/nar/gky1106.

26. Longtine, M.S., McKenzie, A. 3rd, Demarini, D.J., Shah, N.G., Wach, A., Brachat, A., Philippsen, P., and Pringle, J.R. (1998). Additional modules for versatile and economical PCR-based gene deletion and modification in Saccharomyces cerevisiae. Yeast 14, 953–961.

27. Lee, B.H., and Amon, A. (2003). Role of Polo-like kinase CDC5 in programming meiosis I chromosome segregation. Science (New York, N.Y.) 300, 482–486.

28. Benjamin, K.R., Zhang, C., Shokat, K.M., and Herskowitz, I. (2003). Control of landmark events in meiosis by the CDK Cdc28 and the meiosis-specific kinase Ime2. Genes & development 17, 1524–1539.

29. Toth, A., Rabitsch, K.P., Galova, M., Schleiffer, A., Buonomo, S.B., and Nasmyth, K. (2000). Functional genomics identifies monopolin: a kinetochore protein required for segregation of homologs during meiosis i. Cell 103, 1155–1168.

30. Matos, J., Lipp, J.J., Bogdanova, A., Guillot, S., Okaz, E., Junqueira, M., Shevchenko, A., and Zachariae, W. (2008). Dbf4-dependent CDC7 kinase links DNA replication to the segregation of homologous chromosomes in meiosis I. Cell 135, 662–678.

31. Tanaka, T., Cosma, M.P., Wirth, K., and Nasmyth, K. (1999). Identification of cohesin association sites at centromeres and along chromosome arms. Cell 98, 847–858.

32. Carlile, T.M., and Amon, A. (2008). Meiosis I is established through division-specific translational control of a cyclin. Cell 133, 280–291.

33. Wiśniewski, J.R., Zougman, A., Nagaraj, N., and Mann, M. (2009). Universal sample preparation method for proteome analysis. Nat Methods 6, 359–362. 10.1038/nmeth.1322.

34. Olsen, J.V., Macek, B., Lange, O., Makarov, A., Horning, S., and Mann, M. (2007). Higher-energy C-trap dissociation for peptide modification analysis. Nat Methods 4, 709– 712.

35. Cox, J., and Mann, M. (2008). MaxQuant enables high peptide identification rates, individualized p.p.b.-range mass accuracies and proteome-wide protein quantification. Nat Biotechnol 26, 1367–1372.

36. Cox, J., Neuhauser, N., Michalski, A., Scheltema, R.A., Olsen, J.V., and Mann, M. (2011). Andromeda: a peptide search engine integrated into the MaxQuant environment. Journal of proteome research 10, 1794–1805.

37. Cox, J., Hein, M.Y., Luber, C.A., Paron, I., Nagaraj, N., and Mann, M. (2014). Accurate proteome-wide label-free quantification by delayed normalization and maximal peptide ratio extraction, termed MaxLFQ. Molecular and Cellular Proteomics 13, 2513–2526. 10.1074/mcp.M113.031591.

38. Schwanhäusser, B., Busse, D., Li, N., Dittmar, G., Schuchhardt, J., Wolf, J., Chen, W., and Selbach, M. (2011). Global quantification of mammalian gene expression control. Nature 473, 337–342. 10.1038/nature10098.

39. Zhang, X., Smits, A.H., Van Tilburg, G.B.A., Ovaa, H., Huber, W., and Vermeulen, M. (2018). Proteome-wide identification of ubiquitin interactions using UbIA-MS. Nature Protocols 13, 530–550. 10.1038/nprot.2017.147.

